# A PCTAIRE family kinase regulates eye and brain size in freshwater planarians

**DOI:** 10.64898/2026.05.24.727569

**Authors:** A. Guixeras-Fontana, A. Ginés, M.D. Molina, F. Cebrià

## Abstract

**Background:** During embryonic development and regeneration, the growth of any organ must be tightly regulated in order to achieve their optimal final size and become fully functional. Freshwater planarians, with their remarkable plasticity and ability to regenerate any part of their body based upon the presence of adult pluripotent stem cells, provide an ideal model to study how the final organ size is regulated during this process. Also, the fact that planarians are constantly growing and degrowing depending on culture conditions, allows us to study how the size of the different organs is determined under homeostatic cell renewal.

**Results:** Here, we investigate the role of *Smed-pctk-1*, a cyclin dependent kinase that belongs to the PCTAIRE subfamily of CDKs, which remains largely understudied. Functional analyses show that *Smed-pctk-1* silencing disrupts the normal size of the cephalic ganglia and results in an overgrowth of the eyes both in homeostatic and regenerating planarians. The increase in eye size correlates to an increase in the number of both progenitor and differentiated eye cell types. Phototaxis behavioral assays reveal that *Smed-pctk-1* RNAi planarians exhibit a precocious sensitivity to light.

**Conclusions:** Overall, our findings identify *Smed-pctk-1* as a key regulator of eye and neural size in planarians, highlighting its contribution to the mechanisms that control organ growth during both regeneration and homeostasis.

## Background

Organ size regulation is a fundamental aspect of development, ensuring that tissues and organs reach appropriate proportions for their proper function. Several studies have addressed this question in order to understand growth and size control during development [1–8].

Planarians provide an excellent model to study organ size regulation because of their remarkable regenerative abilities. These freshwater flatworms can regenerate any part of their body thanks to the presence of adult pluripotent stem cells, called neoblasts [9–11]. Following an injury, planarians rapidly close the wound and generate a structure named blastema where the neoblasts differentiate into all the missing structures in a two-week process [12]. Moreover, even intact, non-regenerating planarians exhibit a high degree of plasticity, undergoing growth or degrowth depending on culture conditions [13–14]. Both in regeneration and homeostasis in intact animals, proper body proportions and axial polarities are re-established and maintained, respectively [15–16]. All these characteristics make planarians an ideal model to study how final organ size is regulated during regeneration and tissue homeostasis.

Among the organs of planarians, the eye is one of the best characterized at the cellular and molecular level [17–19]. A current model proposes that pluripotent neoblasts give rise to specialized eye progenitors that co-express the transcription factors *ovo, six1*/2, *eya* [20–22]. These specialized progenitors subsequently produce the two eye cell types: sensory photoreceptor neurons and pigmented eye-cup cells [23]. Later in differentiation, photoreceptor neurons become further specified by *otxA* and *soxB* expression [24], while pigmented eye-cup cells are specified by *egfr*-1, *sp6*-9 and *dlx* [25,26]. Several studies have shown that silencing certain genes leads to a reduction or complete absence of the eyes, indicating that these genes promote eye development [20,21,22,24,25,27,28]. Other studies have identified the role of specific genes in correct body patterning and the prevention of ectopic eye formation [19]. However, up to date, few studies have reported factors that act restricting eye. One of the few examples is *Smed-bmp*, whose inhibition leads to an expansion of the photoreceptor cells population [29].

In this study, we investigated the function of *Smed-pctk-1*, a cyclin-dependent kinase that belongs to the PCTAIRE subfamily of CDKs (Cyclin Dependent Kinases), which remains poorly characterized. PCTAIRE kinases (PCTKs) are highly conserved serine/threonine kinases closely related to canonical CDKs. PCTKs subfamily differ from other canonical CDKs at structural level for the substitution of the conserved serine residue in the canonical PSTAIRE motif by a cysteine. This modification, absent in the early characterized CDKs, has been proposed to alter cyclin binding and create a unique interface for alternative cofactors [30,31]. Unlike canonical CDKs, PCTKs are often enriched in post-mitotic neurons of adult mammalian brains, suggesting important neuron-specific functions independent of cell cycle regulation [32,33]. Given their expression in specialized tissues such as brain, testis, post-mitotic neurons, and elongated spermatids, research has primarily explored their potential contributions to cellular differentiation and tissue-specific processes [30,34]. Therefore, in vivo knockdown experiments in model organisms such as planarians can provide a valuable opportunity to uncover their roles in development and tissue differentiation in the context of regeneration and homeostatic cell turnover.

Here we identified *Smed-pctk-1* as a negative regulator of eye growth in both homeostatic and regenerating planarians. The silencing of *Smed-pctk-1* impacts eye progenitor dynamics and results in changes in the number of differentiated photoreceptor neurons and eye pigmented cup cells. Additionally, its silencing results in enhanced phototaxis responses to light and smaller regenerated brains. Overall, our results reveal a previously uncharacterized role for PCTKs in regulating organ size in regenerative systems.

## Results

### *Smed-pctk-1* is mainly expressed in the central nervous system

The identified *Smed-pctk-1* clustered together, in our phylogenetic analysis, with other CDKs from the CDK16/17/18 subfamily carrying within it the characteristic PCTAIRE domain of this non-canonical CDKs subfamily (Additional file 1: Fig. S1). To investigate its expression pattern, we performed whole-mount *in situ* hybridization (WISH) in both intact and regenerating animals (Fig. 1). In intact animals, *pctk-1* expression was detected throughout the planarian body, with a distinctly stronger signal in the central nervous system (CNS), which in planarians consists of a bilobed cephalic ganglia and two ventral nerve cords (Fig. 1A). The expression pattern of *pctk-1* through the mesenchyme does not correspond with neoblasts population, as its expression is not affected after irradiation, treatment that specifically eliminates neoblasts [35] (Additional file 2: Fig. S2). Moreover, *Smed-pctk-1* was also detected in the photoreceptors (Fig. 1B). During anterior regeneration, *pctk-1* expression was clearly detected in the newly formed cephalic ganglia (cg) from early stages of regeneration (Fig. 1C, arrowheads)

**Fig. 1.**
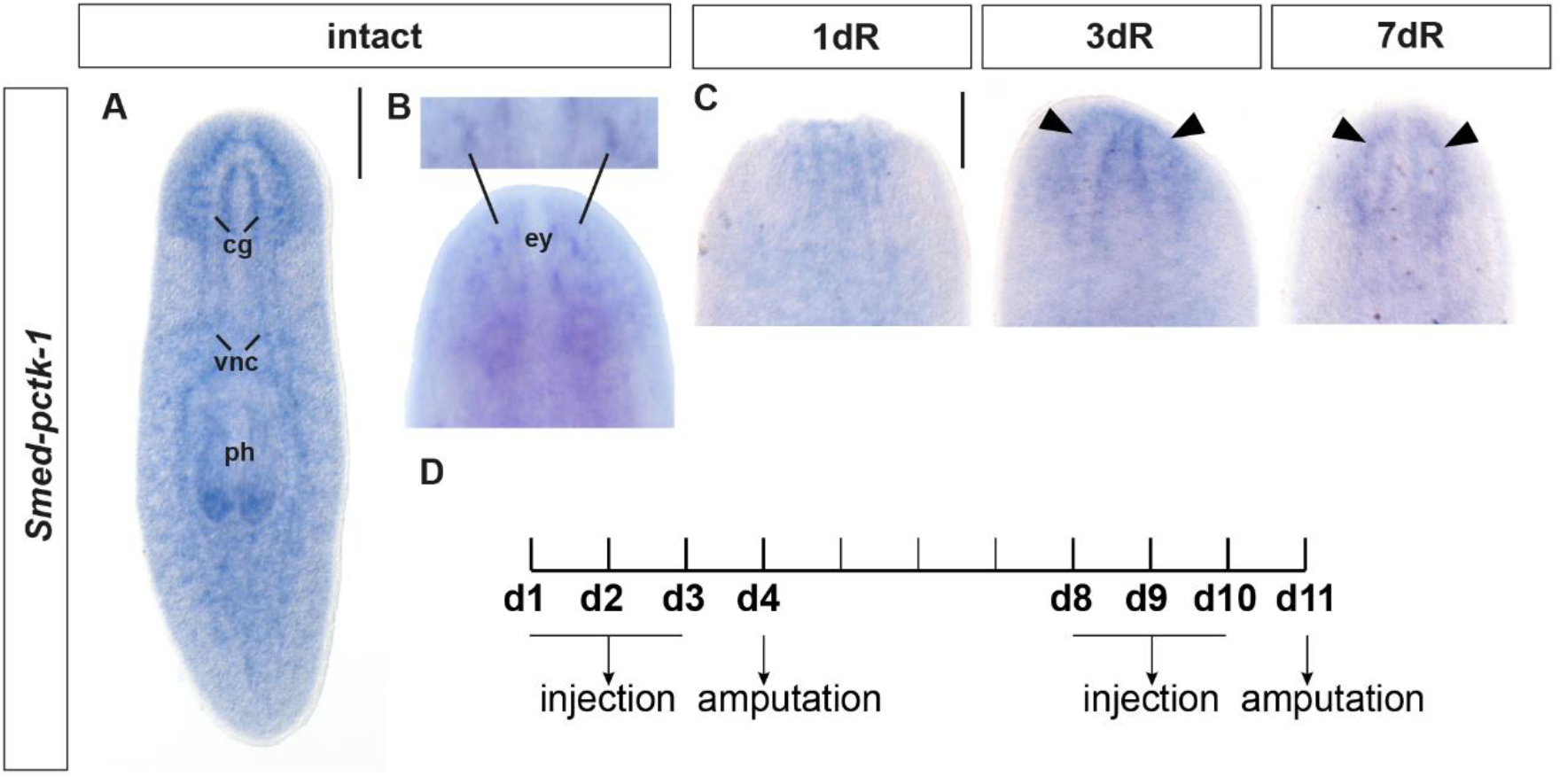
*Smed-pctk-1* expression in intact and regenerating planarians. **A**,**B** *Smed-pctk-1* expression pattern detected by whole mount *in situ* hybridization (WISH) in intact animals, ventral (A) or dorsal (B) views. cg, cephalic ganglia; vnc, ventral nerve cords; ph, pharynx; ey, eyes. **C** *Smed-pctk-1* expression pattern at 1, 3 and 7 days of regeneration (dR). Arrowheads point to the cephalic ganglia. **D** Schematic representation of the RNAi experimental procedure. Animals went through two rounds of dsRNA injection and amputation. Scale bars: 400 μm.

### *Smed-pctk-1* is required for proper cephalic ganglia growth

As *Smed-pctk-1* was mainly enriched in the CNS we aimed to analyze whether its silencing could affect the regeneration of the cephalic ganglia. Therefore, we silenced the expression of *Smed-pctk-1* by RNA interference (RNAi) and assessed its impact on regenerating animals (Fig. 1D). Whole mount *in situ* hybridization was performed using specific markers for neural progenitors and differentiated neurons (Fig. 2). Neural progenitor cells were visualized with *Smed-sim* [36] (Fig. 2A,D), whereas mature brain lateral branches were labelled with *Smed-gpas* [37] (Fig. 2G). Notably, *Smed-sim* labels neural progenitors as well as differentiated cells [36]. We quantified the number of progenitor *sim*+ cells, normally located between the two cephalic ganglia (white dashed triangle). We observed a significant decrease in the number of *sim*+ progenitor cells in *pctk-1* silenced animals at 5 days of regeneration compared to controls (Fig. 2B). Despite this reduction, the length of the newly regenerated cephalic ganglia (black dashed circle), did not differ significantly at this stage (Fig. 2C). However, by 13 days of regeneration (Fig. 2D), the effects of *pctk-1* silencing became more pronounced. We observed a further decrease in the number of *sim*+ progenitor cells, located in the medial region of the planarian body at this point (white dashed box) (Fig. 2E), and the cephalic ganglia were smaller compared to controls (Fig. 2F). Using the *Smed-gpas* neural marker, which labels the lateral branches, we confirmed that the cephalic ganglia regenerated after *Smed-pctk-1* silencing were smaller than in controls, with quantitative analysis of brain width, length and area showing a significant reduction (Fig. 2G-J). Given that canonical CDKs are key regulators of the cell cycle, and despite that the non-canonical subfamily of PCTAIRE CDKs seem to have no role in cell cycle, we evaluated whether silencing *pctk-1* altered the pool of neoblasts and/or their proliferative activity. Neoblasts are defined by the expression of *piwi* whereas proliferating neoblasts can be detected using an anti-PH3 antibody, which marks cells in the G2/M phase [38,39]. No significant differences were observed either in the total neoblast pool that expresses *piwi* or the proliferating subset (Additional file 3: Fig. S3), suggesting that *pctk-1* would not affect neoblast maintenance and proliferation. Together, these results indicate that *Smed-pctk-1* would be necessary for proper cephalic ganglia growth, likely by regulating neural progenitor dynamics rather than overall stem cell proliferation.

**Fig. 2.**
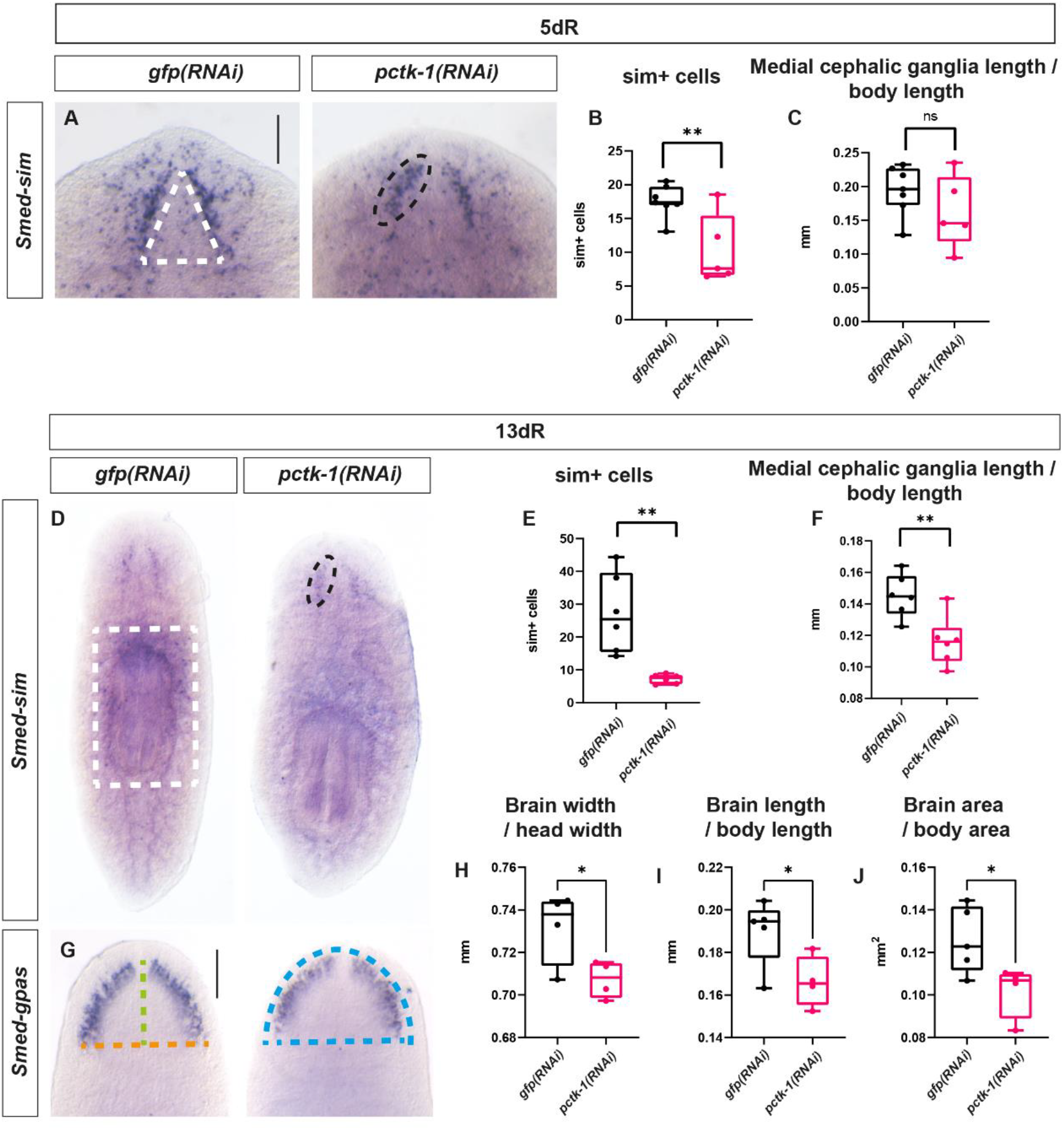
Nervous system defects after *Smed-pctk-1* silencing. **A**,**D** Neural progenitor cells labelled by WISH for *Smed-sim*. **B**,**E** Quantification of neural progenitor cells (*sim*+ cells) within the white boxed areas. **C**,**F** Quantification of medial cephalic ganglia length within black boxed areas (^ns^p-value > 0.05,**p-value < 0.01, Student’s t-test). Values represent the mean of at least 5 animals per condition. **G** Brain lateral branches labelled by WISH for *Smed-gpas*. **H-J** Quantification of brain width (orange dashed line), length (green dashed line) and area (blue boxed area) (*p-value < 0.05, Student’s t-test). Values represent the mean of at least 4 animals per condition. Scale bars: 200 μm. Images at 5 and 13 days of regeneration (dR).

### *Smed-pctk-1* restricts eyes size during regeneration

During anterior regeneration *Smed-pctk-1* silenced animals displayed normal-sized blastemas (Fig. 3A). Externally, the first organs that become visible within the regenerative blastema are the pigmented eye-cups (around 3-4 days of regeneration). The new pigmented eye-cups became clearly visible after 5 days of regeneration (black arrowheads). At this early stage, no significant differences were observed in the area of the pigmented eye-cup relative to the body area between controls and *pctk-1* RNAi planarians. However, by 12 days of regeneration, a significant increase in the pigmented eye-cup area was detected in *pctk-1* silenced planarians (Fig. 3A,B). From day 12 onward, the entire newly formed eye, including both the pigmented-eye cup and the white region (white arrowheads) where the photosensitive cells reside, is distinguishable and quantifiable. By 24 days of regeneration, a significant increase in the total area of the eye was observed (Fig. 3A,C). This enlargement in both the pigmented eye-cup area and the total eye area persisted from 24 to 53 days after amputation, showing the long-term effects of *pctk-1* silencing (Fig. 3).

**Fig. 3.**
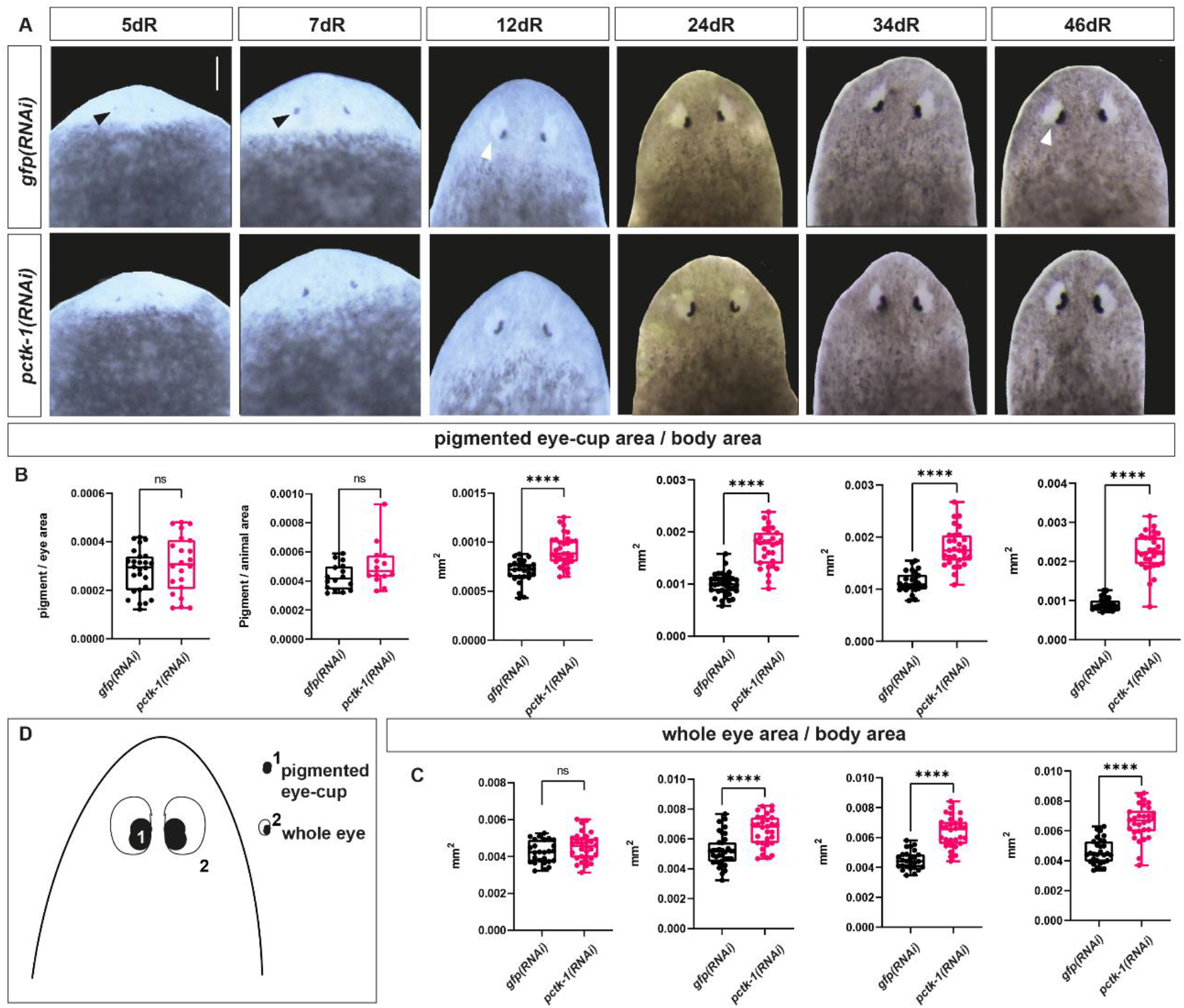
*Smed-pctk-1* silencing results in bigger eyes. **A** In live animals both the whole eye region (white arrowheads) and the pigmented eye-cup region (black arrowheads) are bigger after *Smed-pctk-1*RNAi animals. Animals at 5, 7, 12, 24, 34, 46 days after amputation (dR). Scale bar: 300 μm. **B** Ratio between the pigmented eye-cup area and the body area. **C** Ratio between the whole eye area and the body area. (^ns^p-value > 0.05; ****p-value < 0.0001, Student’s t-test). Values represent the mean of at least 12 eyes per condition. **D** Schematic representation of pigmented ^1^eye-cup and ^2^whole eye regions.

### Smed-*pctk-1* silencing results in an increase of both progenitor and differentiated eye cells

As described previously, planarian eyes are constituted by two cell populations, pigmented cells that form the eye-cup and photoreceptor neurons, whose cell bodies cluster around the pigmented eye-cups whereas their axons project to the cephalic ganglia [17]. These two cell populations derive from a common progenitor [24]. Given that eyes appeared larger in *Smed-pctk-1* live animals, we assessed the effect of *Smed-pctk-1* silencing on these specific eye cell populations using whole-mount *in situ* hybridizations with specific markers for both progenitor and differentiated eye cells (Fig. 4). Eye progenitor cells were labelled with *Smed-ovo-1*, which is expressed in both progenitors and differentiated eye cells; being eye progenitors distributed in two rows of cells posterior to the mature eyes [24]. On the other hand, differentiated pigmented eye-cup cells were specifically labeled with *Smed-tph* [40], and differentiated photoreceptor neurons were visualized with VC-1, an antibody specific for arrestin [17] (Fig. 4A). In regenerating *pctk-1-*silenced animals, we observed a significant increase in the number of *ovo1*+ progenitor cells at 5 and 13 days of regeneration. By 53 days of regeneration, when animals are fully regenerated and only homeostatic processes are active, *Smed-pctk-1 RNAi* animals tended to present higher numbers of progenitors compared to controls, although no significant differences in progenitor numbers were detected (Fig. 4B). Regarding differentiated eye cell populations, the number of photoreceptor neurons at 5 days of regeneration showed no significant change (Fig. 4C). However, at 13 and 53 days after amputation, the number of both differentiated pigmented eye-cup cells and photoreceptor neurons was significantly increased in *pctk-1* RNAi animals (Fig. 4D), suggesting that the early expansion of progenitor cells drives a subsequent increase in the mature eye cell populations, leading to larger eyes.

**Fig. 4.**
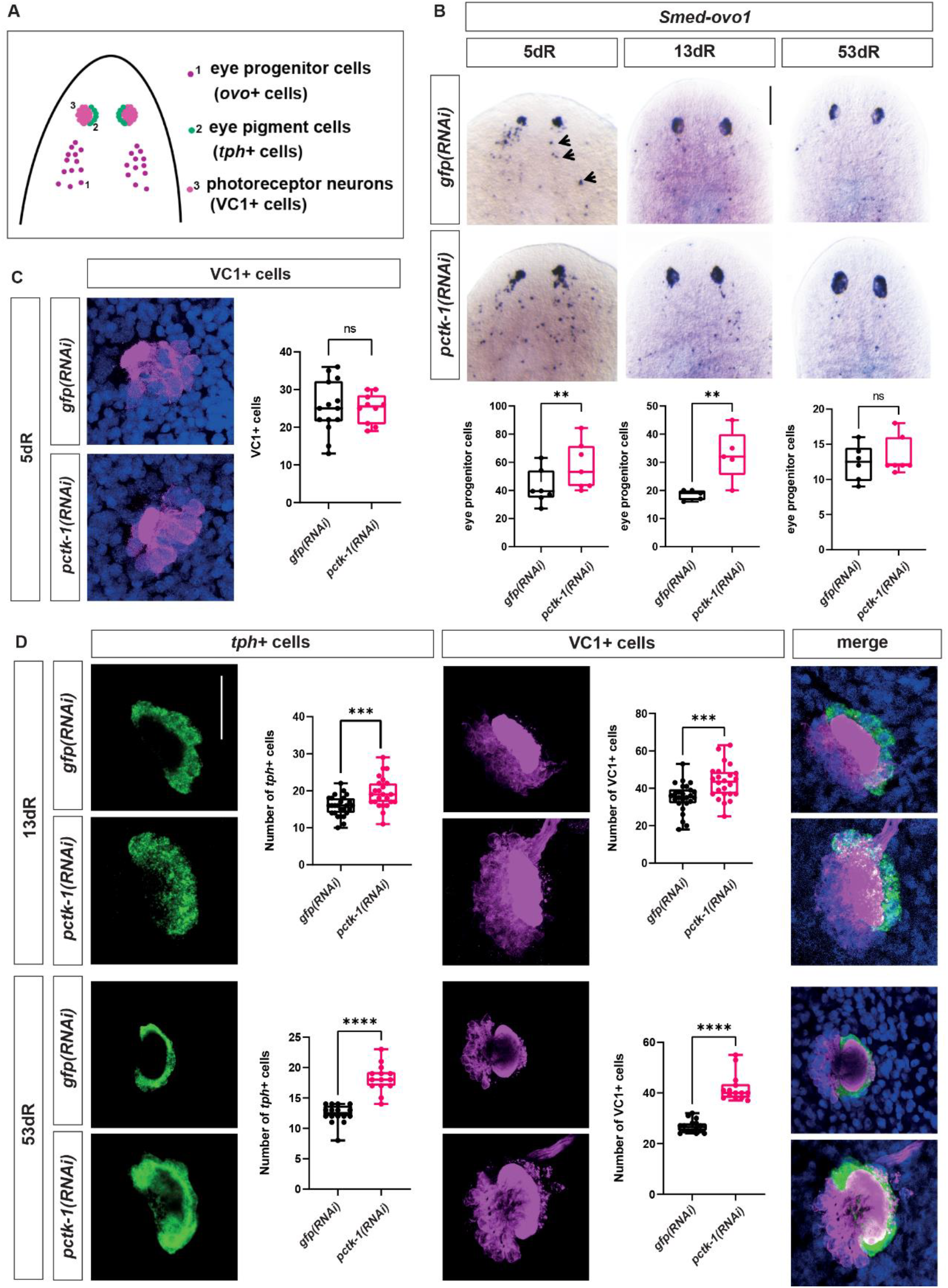
Increase in eye progenitor and mature cell types after *Smed-pctk-1* silencing. **A** Schematic representation of ^1^progenitor eye cells (*ovo+* cells), ^2^pigment-cup cells (*tph+* cells) and ^3^photoreceptor neurons (VC1+ cells). **B** Eye mature and progenitor cells labelled by WISH for *Smed-ovo-1*. Quantification of eye progenitor cells (black arrows; ^ns^p-value > 0.05; **p-value < 0.01, Student’s t-test). Values represent the mean of at least 5 animals per condition. **C** Immunostaining with VC-1 (anti-arrestin) antibody specific for photoreceptor cells (magenta) and quantification of VC1+ cells. Cell nuclei labelled with DAPI. Quantification of VC1+ cells (^ns^p-value > 0.05, Student’s t-test). **D** Double FISH and immunostaining and quantification of *tph+* (in green) and VC1+ (in magenta) cells. Quantification of *tph+* and VC1+ cells (***p-value < 0.001; ****p-value < 0.0001, Student’s t-test). Values represent the mean of at least 12 eyes per condition. Scale bars: 200 μm in B and 100 μm in C and D. Images at 5, 13 and 53 days after amputation (dR). Cell nuclei labelled with DAPI.

### 3D expansion of photoreceptor neurons after silencing *pctk-1*

In planarians, the visual axons of the photoreceptor neurons project either ipsilaterally to the inner medial region of the cephalic ganglia, or contralaterally forming an optic chiasm [17,41]. The cell bodies of the planarian photoreceptor cells are placed more externally and dorsal relative to the cephalic ganglia, whereas their axons project to inner, more ventral regions; the optic chiasm is formed along the commissure that connects both cephalic ganglia [12,17]. Confocal microscopy allowed us to visualize how, in control planarians, the cell bodies of the photoreceptor neurons and their axonal projections are spatially separated in more outer (dorsal) and inner (ventral) regions, respectively (Fig. 5A). In contrast, after silencing *Smed-pctk-1*, the increased number of photoreceptor neurons described above (Fig. 4) resulted in cell bodies being located abnormally in more inner regions (Figs. 5A and Additional file 4: Fig. S4). In addition, the optic chiasm appeared thicker in these animals (arrowheads), likely reflecting a higher number of axonal projections due to the expanded photoreceptor neuron population (Fig. 5A). Finally, measurements revealed that the length of the visual axons (as a whole) from the chiasm towards posterior was also significantly increased after silencing *pctk-1* (Fig. 5B-C). That is, the visual axons (considered altogether) projected to more posterior regions of the cephalic ganglia.

**Fig. 5.**
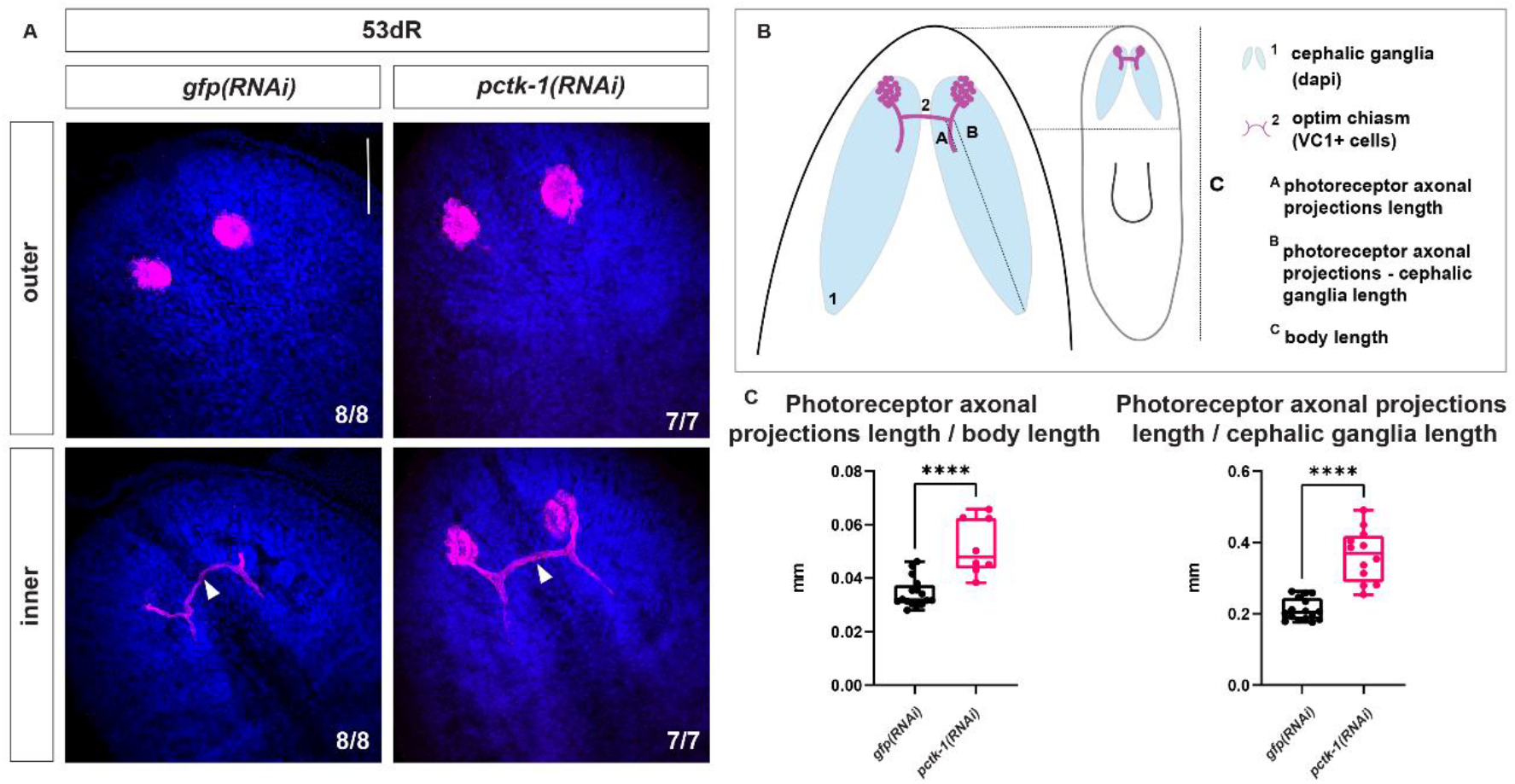
Ventral expansion of photoreceptor neurons after *Smed-pctk-1* silencing. **A** Photoreceptor neurons and optic chiasm visualized by VC-1 immunostaining. Z-projection of the more outer (dorsal) or inner (ventral) confocal sections of the region where the eye cells (outer) and the visual axons and chiasm (inner) are located. **B** Schematic representation of ^1^cephalic ganglia (DAPI*+* cells) and ^2^photoreceptor neurons and optic chiasm (VC1+ cells in magenta). **C** Ratio between the length of the photoreceptor axonal projections and the length of the body, and ratio between the length of the photoreceptor axonal projections and the length of the cephalic ganglia (***p-value < 0.0001, Student’s t-test). The length of the axonal projections of the photoreceptors was measured from the chiasm to their most posterior end. Values represent the mean of at least 12 eyes per condition. Scale bars: 200 μm. Images at 53 days post amputation (dR).

### *Smed-pctk-1* RNAi planarians exhibit an increased sensitivity to light

Planarians typically display negative phototaxis, moving away from light when exposed to it [18]. To determine whether the increase in eye size and number of photoreceptor neurons observed after *Smed-pctk-1* silencing affects this behavior, we performed phototaxis behavioral assays in regenerating planarians (Fig. 6). Animals were exposed to a light gradient and their movement over time was recorded (Fig. 6A). We first quantified the percentage of animals located in different regions away from the direct light source at different time points (Fig. 6B). The results showed that, at 7 days of regeneration, *pctk-1* silenced planarians moved slightly faster to darker regions compared to controls. Also, the total distance traveled by *pctk-1* silenced planarians was significantly longer, at the two recorded time points (t1 and tf) (Fig. 6C). No significant differences were detected at later stages (data not shown). To better distinguish possible differences between treatments, we additionally analyzed regenerating tails. As described in regenerating trunks (Fig. 3), the pigmented eye-cup area relative to body area was significantly larger in *Smed-pctk-1* silenced tails at 7 days of regeneration. Consistent with trunk results, *pctk-1* silenced regenerating tails also tended to move faster to darker regions and traveled farther away from the light source than controls, indicating an enhanced phototactic response to light in both regenerating fragments (Additional file 5: Fig. S5).

**Fig. 6.**
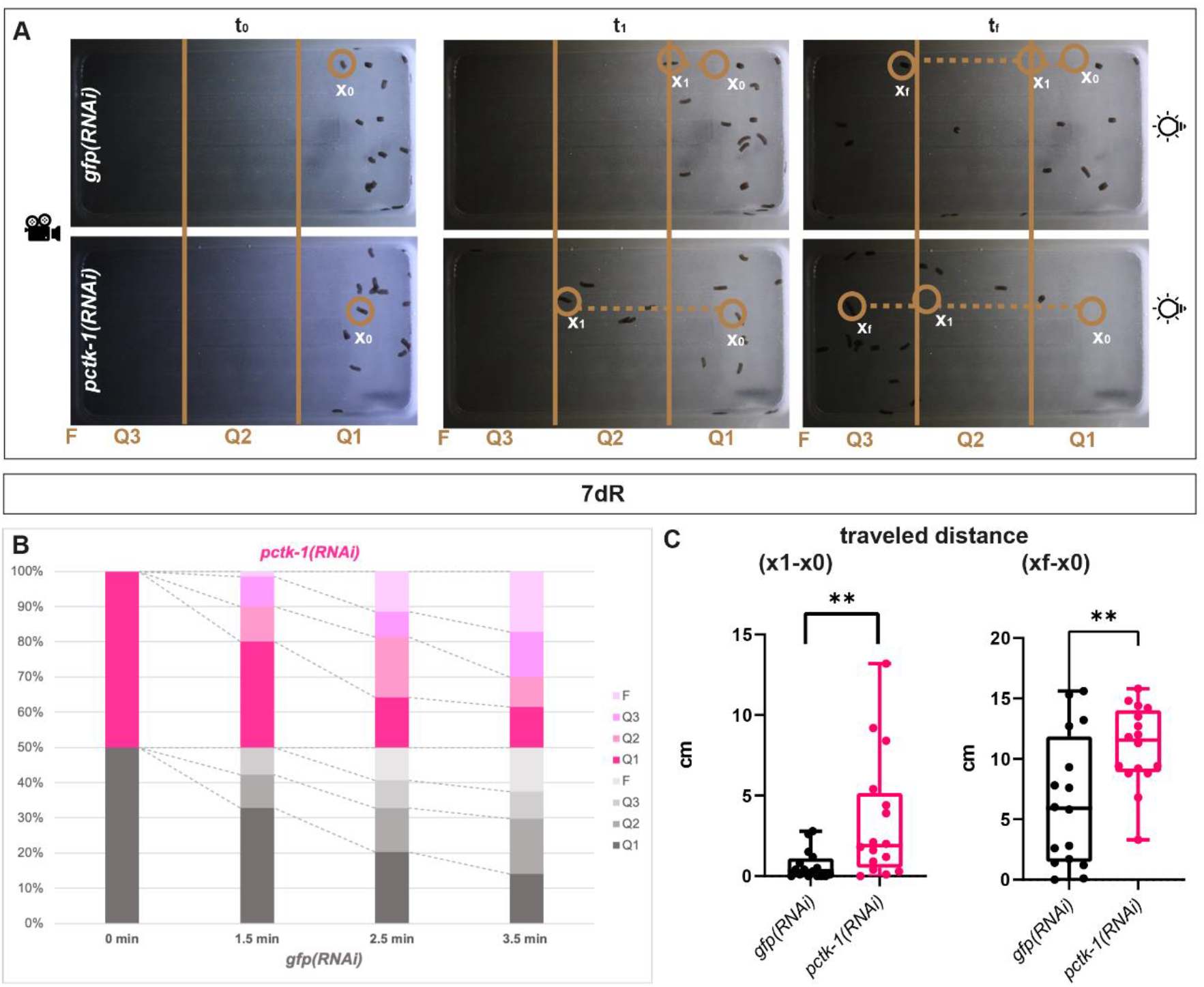
Increased light sensitivity after *Smed-pctk-1* silencing. **A** Representative image of the phototaxis behavioral assay. **B** Light responses measurements by quantifying the percentage of animals in Q1, Q2, Q3 and F regions at various time points after the exposition of direct light initial t0 (0 mins), intermediate t1 (1 min) and final tf (3,5 mins). **C** Light responses measurements by calculating the distance in cm (X1-X0) and (Xf-X0) after the exposition to direct light. (**p-value < 0.01, Student’s t-test). Values represent the mean of at least 16 animals per condition. Animals at 7 days of regeneration (dR).

### *Smed-pctk-1* also regulates eye size in homeostasis

Having observed the effects of *Smed-pctk-1* during regeneration, we next asked whether similar effects occurred in intact, non-regenerating animals. Indeed, we found that silencing *pctk-1* also led to an increase in the eye size in non-regenerating animals (Fig. 7). At 7 days post *Smed-pctk-1* silencing the pigmented eye-cup area was significantly larger (Fig. 7A,C,E), and after 6 weeks, both the pigmented eye-cup area and the total eye area were increased (Fig. 7B,D,F). Overall, *Smed-pctk-1* seems to be regulating eye final size both during regeneration and homeostasis.

**Fig. 7.**
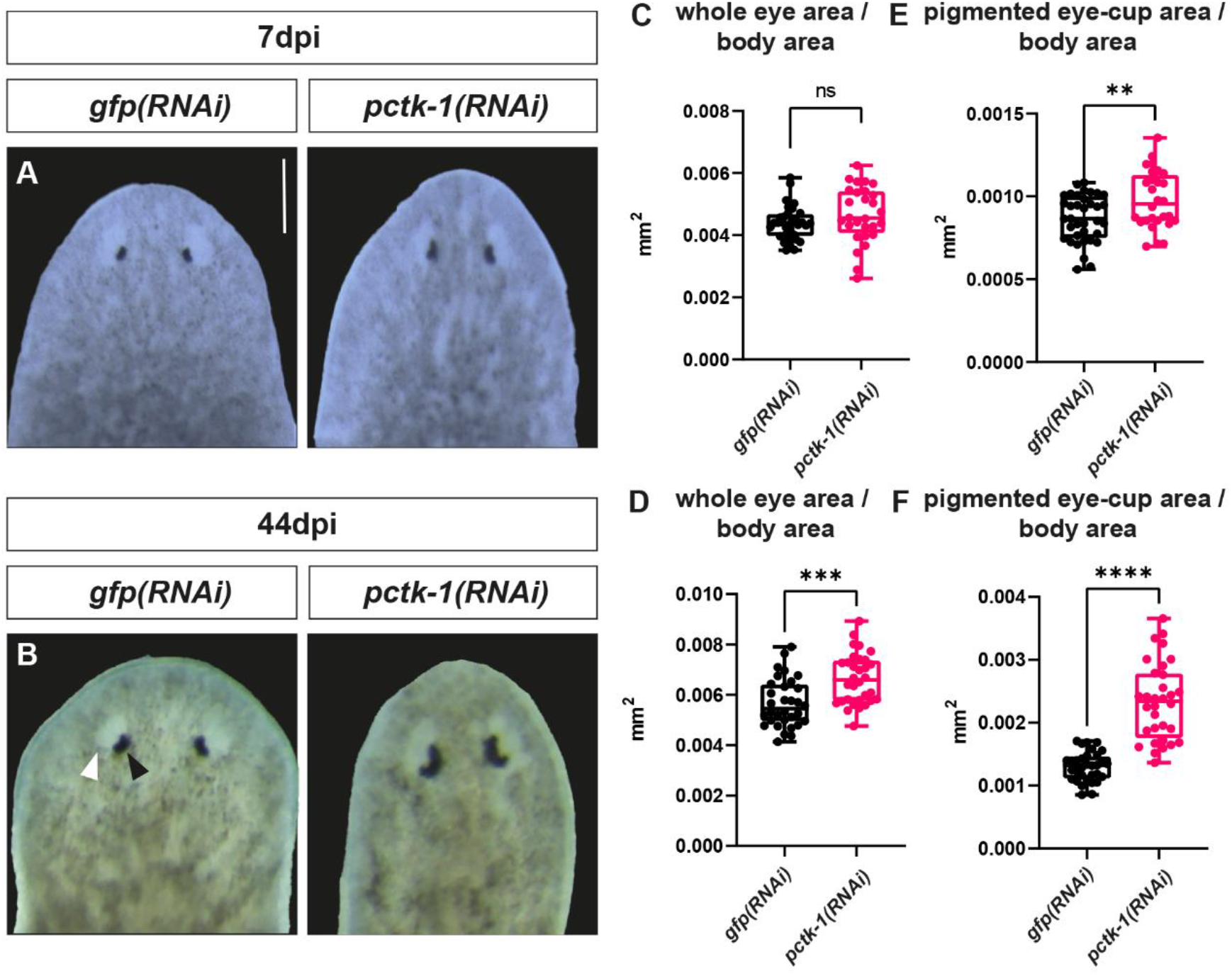
*Smed-pctk-1* silencing results in bigger eyes in intact homeostatic planarians. **A, B** Live intact animals show that both the whole eye region (white arrowheads) and the pigmented eye-cup region (black arrowhead) are bigger in *Smed-pctk-1* RNAi animals through time in homeostatic conditions. Animals at 7 and 44 days post injection (dpi). Scale bar: 300 μm. **C**,**D** Ratio between the pigmented eye-cup area and the total body area. **E**,**F** Ratio between the whole eye area and the total body area. (**p-value < 0.01; ****p-value < 0.0001, Student’s t-test). Values represent the mean of at least 30 eyes per condition.

## Discussion

Understanding the gene regulatory mechanisms that govern stem cell proliferation and differentiation is a central question in developmental biology. Freshwater planarians provide an ideal model system due to their remarkable ability to regenerate any part of their body based upon the presence of adult pluripotent stem cells [11,42,43]. Although many studies have uncovered the role of several genes and signaling pathways in the regeneration of specific organs and tissues such as the brain [38,44,45], muscles [46], eyes [25, 27], epidermis [47,48], excretory [49] or the digestive system [50,51], much little is known about how the final size of these organs is achieved. A previous study reported that silencing BMP4 in regenerating planarians results in an increase of eye pigment cells and photoreceptor neurons together with a clear patterning defect of the pigment cells and the visual axonal projections [29]. Here, we have identified *Smed-pctk-1* as a gene required for regulating the final size of the planarian eyes in both regeneration and homeostatic contexts. In contrast to the aforementioned study, *Smed-pctk-1* silencing does not cause major patterning defects.

Protein kinases are central regulators of cellular signaling pathways, and their dysregulation is implicated in diverse pathologies, particularly cancer [30]. However, the biological roles of many kinases, including a substantial fraction of cyclin-dependent kinases (CDKs), remain poorly defined. CDKs, once considered primarily cell cycle regulators [52], are now recognized as multifunctional enzymes involved in DNA replication, repair, transcription, and neuronal processes. Evolutionary diversification has expanded the CDK family from six members in yeast to twenty-one in humans, with atypical CDKs such as CDK5 and the PCTAIRE subgroup (CDK16–18) acquiring specialized functions [53]. Within this subgroup, PCTKs are particularly intriguing: they are expressed in differentiated tissues, especially in testis, skeletal muscle, brain and neural structures [32,34,54–60], yet their activation mechanisms and physiological roles remain largely unexplored. Some studies have shown a role of PCTKs in the control of neurite outgrowth [32], dendrite development [58], skeletal myogenesis [59] or oligodendrocytes differentiation [60]. Recent knockout studies in mice further suggest developmental and potentially sex-specific functions, but published data are scarce [30,61].

Consistent with the observations made in other models [32,34,54–60], in planarians *Smed*-*pctk-1* is expressed in postmitotic differentiated cells throughout the planarian body, specifically in neurons and photoreceptor cells (Fig. 1). In agreement with the suggested roles for the genes of the PCTAIRE subgroup, *Smed-pctk-1* silencing does not alter neoblast abundance or affect cell proliferation, suggesting that it does not regulate cell cycle (Additional file 1,3:, Fig. S1,S3). Remarkably, however, our studies uncover the role of *Smed-pctk-1* in regulating organ size by affecting the number of progenitors during both regeneration and homeostasis in non-regenerating animals. Thus, its silencing results in smaller brains which correlates with a decrease in the number of neural progenitors (Fig. 2). On the other hand, the silencing of *Smed-pctk-1* impairs eye regeneration in the opposite direction to that described for the brain, as bigger eyes differentiate in both regenerating and non-regenerating animals as a result of an increase in the number of specialized progenitors (Fig 3, 7). Despite these opposite outcomes for brain and eye fate, in both cases the effects on final organ size seem to be mediated by the regulation of the number of specialized progenitors, consistent with previous works showing that eye progenitor abundance is a key determinant for eye regeneration and homeostasis [20,24,25]. *Smed-pctk-1* may independently regulate brain and eye progenitors in opposite directions, or alternatively, may influence fate decisions from a yet uncharacterized and hypothetical common progenitor population for eye and brain cells, as suggested by recent lineage tracing and single cell studies for other planarian cell types, i.e. muscle, neural and secretory cells [62–64]. Even though our *in situ* hybridizations have not been able to detect *Smed-pctk-1* expression neither in neoblasts nor in eye and neural progenitors, we cannot exclude the possibility that this gene is express either at very low levels or in a small subset of these cell populations. Additional experiments would be required to determine whether the effects observed here are autonomous or non-autonomous, for instance, potentially mediated through the muscle cells, which have been shown to be an important source of signals to control cell specification and differentiation in planarians [64–67].

In the case of the eyes, the expansion of eye progenitor cells during early regeneration indicates that *Smed-pctk-1* normally acts to limit progenitor eye population during eye formation. After full regeneration is accomplished, *Smed-pctk-1-RNAi* animals retain a higher number of differentiated eye cells compared to controls (Fig. 4). However, no differences in the number of eye progenitors are detected at this stage. This could be due to the strong reduction of eye progenitors production associated with a prolonged period of starvation, even in control animals, which might difficult detecting any statistically significant difference between treatments [68]. At the same time, the higher number of differentiated eye-pigment and photoreceptor cells is maintained high even at these late stages, most probably as a result of their accumulation all along the duration of the experiment. In any case, the persistence of an increased number of differentiated cell numbers at 53 days post-amputation, when regeneration-associated proliferation has ceased, indicates that *Smed-pctk-1* contributes to long-term tissue size maintenance rather than transient regenerative responses.

The cell bodies of the photoreceptor neurons are normally placed dorsal to the cephalic ganglia whereas their axonal projections project ventrally in two trajectories: some of them project contralaterally along the posterior side of the commissure connecting both cephalic ganglia and forming an optic chiasm, and others project ipsilaterally along the medial region of the cephalic ganglia [41,69]. Our results show that the increased number of photoreceptor cells results in their cell bodies occupying more ventral positions, closer to the cephalic ganglia (Fig. 5 and additional file 4: Fig. S4). Their axonal projections, however, follow normal trajectories, indicating that *Smed-pctk-1* does not affect the direction of the growth of the visual axons. Nevertheless, it is important to note that in *Smed*-*pctk-1-RNAi* animals the visual axons, considered altogether, extend to more posterior regions respect to the cephalic ganglia. This is not only due to the fact that, at the same time, those cephalic ganglia are shorter, as the total length of the axonal projections from the chiasm to their most posterior end is also longer when normalized to the body length. Due to the current impossibility to measure the length of visual axons from individual photoreceptor cells, it remains unclear whether *Smed-pctk-1* silencing generate longer axons that project to more posterior regions of the brain or, alternatively, whether the excess of photoreceptor cell bodies located closer to the cephalic ganglia leads to an apparent posterior extension without changes in individual axon length.

Finally, as planarians show negative phototaxis, we examined whether the increase in photoreceptor neurons had an impact on this behavior (Fig. 6 and additional file 5: Fig. S5). At 7 days of regeneration, *Smed-pctk-1-silenced* planarians show an enhanced phototactic response and sensitivity to light, as they move further from the light source compared to controls. This would agree with the observation of those animals having a larger eye-pigment area compared to controls. Importantly, the maintenance of a coordinated and directional escape response indicates that the core neural circuits underlying phototaxis remain functional despite changes in photoreceptor cell number and cephalic ganglia size. On the other hand, in fully regenerated animals, no differences in the phototaxis response are observed despite the persistently higher number of photoreceptor cells (data not shown). This may reflect that a fraction of the excessive photoreceptor cells is not functional (which is improbable as at early stages of regeneration those cells function properly to trigger a phototactic response), or more probable, a saturation effect in which an increase beyond a certain number of photoreceptor cells does not further enhance the phototactic response.

In summary, we have identified *Smed-pctk-1* as a key regulator of organ size in both regenerative and homeostatic contexts.

## Conclusions

Our study identifies *Smed-pctk-1* as a previously uncharacterized regulator of organ size in planarians, acting during both regeneration and homeostasis. Unlike canonical CDKs, *Smed-pctk-1* does not affect neoblast maintenance or global cell division but instead modulates the number of lineage-specific progenitors, controlling the final size of distinct organs. Notably, *Smed-pctk-1* exerts opposite effects on different tissues, restricting eye progenitor expansion while promoting neural progenitor production, as its silencing leads to bigger eyes and smaller brains. Overall, our work uncovers a novel role for a PCTAIRE kinase in regulating progenitor dynamics and organ growth, expanding the functional repertoire of atypical CDKs beyond their established roles. These findings provide new insight into how tissue size is controlled in regenerative systems and establish planarians as a valuable model to study kinase-mediated regulation of stem cell differentiation and organ homeostasis.

## Methods

### Animal culture

An asexual clonal line of the planarian species *Schmidtea mediterranea* was maintained at 20ºC in 1x PAM water (1.6 mM NaCl, 1.0 mM CaCl2, 1.0 mM MgSO4, 0.1 mM MgCl, 0.1 mM KCl, and 1.2 mM NaHCO3, dissolved in deionised water) at pH 7.0. Planarians were fed once a week with organic veal liver and starved at least one week before any experimental procedure. For knockdown experiments, we selected animals around 6 mm in length. For irradiation experiments, planarians were exposed to 85 grays of X-irradiation using a YXLON SMART 200 irradiator.

### Sequence amplification and knock-down

#### Primary PCR

To amplify *pctk-1* we used cDNA from wild type *S. mediterranea* worms. Primary PCR was performed using 0.5 µL of cDNA, 3.25 µL of 10x Dream Taq Reaction Buffer, 0.5 µL of dNTPs (10 mM), 0.25 µL of Dream Taq DNA Polymerase, 1.25 µL of Primer Forward (2.5 μM), 1.25 µL of Primer Reverse (2.5 μM) and 18 µL of water. The primer sequences were as follows: GGCCGCGGacgaagaggaagccaaggac (*pctk-1*-F), GGCCGCGGacgaagaggaagccaaggac (*pctk-1*-R). *Smed-pctk-1* primers were designed from the PlanMine sequence SMEST023289001.1. For all genes, both primer pairs included linkers for Universal T7/SP6 primers: GGCCGCGG (linker-F) and GGCCGCGG (linker-R). The thermocycler programme used was: 95ºC (30 s); 35 cycles at 95ºC (30 s), 57 ºC (30 s) and 72ºC (60 s); and 72ºC (5 min). We assessed 3 µL/sample of PCR products in a 1% agarose gel and the remaining volume was used for secondary PCR.

#### Secondary PCR

We prepared 100 µL reactions using 2 µL of primary PCR, 13 µL of 10x Dream Taq Reaction Buffer, 2 µL of dNTPs (10 mM), 1 µL of Dream Taq DNA Polymerase, 5 µL of Universal T7-F5’ primer (2.5 µM, GAGAATTCTAATACGACTCACTATAGGGCCGCGG) and 5 µL of Universal T7-R3’ primer (2.5 µM, AGGGATCCTAATACGACTCACTATAGGCCCCGGC) or 5 µL of Universal SP6-R3’ primer (2.5 µM, AGGGATCGATTTAGGTGACACTATAGGGCCCCGGC) and 72 µL of water. Samples were run in a thermocycler as follows: 95ºC (30 s); 35 cycles at 95ºC (30 s), 57 ºC (30 s) and 72ºC (60 s); and 72ºC (5 min). The size of the bands was assessed by running 3 µL/sample in a 1% agarose gel. The remaining volume was purified by QIAquick commercial kit. Purified samples were eluted in 30 µL of nuclease-free water.

#### dsRNA synthesis

For each sample, we mixed 1 µg of purified cDNA, 8 µL of 5x Transcription Buffer (Thermo Scientific), 4 µL of dNTPs (25 mM), 2 µL of RNase Inhibitor (Applied Biosystems by Thermo Fisher Scientific), 4 µL of T7 RNA Polymerase (Thermo Scientific) and up to 40 µL of nuclease-free water and incubated for 4 hours at 37ºC. Then, we added 1 µL of DNase (1 U/µL, Thermo Scientific) and incubated for another 30 min at 37ºC. After incubation, reactions were stopped with 360 µL of Stop Solution (1M NH4OAc, 10 mM EDTA, 0.2% SDS). The resulting dsRNA was purified using phenol:chloroform. We added 1 µL of Glycogen and 400 µL of acid phenol:chloroform (pH 4.5, Thermo Fisher) per reaction and vortexed thoroughly. We centrifuged for 10 min and collected the aqueous top phase into a new tube. We added 400 µL of chloroform, centrifuged for 10 min, and collected the top phase again. We incubated this phase for 20 min at 68ºC and for 45 min at 37ºC for proper annealing. To precipitate pellets, we added 0.5 µL of Glycogen and 1 mL of cold ethanol, vortexed and centrifuged for 20 min. Pellets were washed in 200 µL of 70% ethanol and centrifuged for 10 min. We discarded supernatants and let the pellets dry for 5 min at 37ºC. Pellets were resuspended in 12 µL of nuclease-free water. All centrifugations were performed at 4ºC and maximum speed. As a quality check, we ran 0.5 µL of purified dsRNA in a 1% agarose gel. Finally, we measured the concentration in a Nanodrop and diluted each dsRNA at a final concentration of 1000 ng/µL.

#### RNA interference

For dsRNA injections, we used worms that were 6 mm in length. We used a Nanoject II injector to deliver single *gfp* (control) and *pctk-1* dsRNA, as previously described [70]. Each animal was injected with 0.1 μg of dsRNA for 3 consecutive days (0.3 μg in total) two rounds separated by a 4-day interval. One day after each round of injection, planarians underwent pre-pharyngeal and post-pharyngeal amputation to induce anterior and posterior regeneration and cultured in PAM water being monitored daily until they were fixed for in situ hybridizations or immunostainings.

### Whole mount *in situ* hybridization

Colorimetric whole mount *in situ* hybridization (WISH) was performed as previously described by [71]. Animals were euthanized by immersion in 5% N-acetyl-L-cysteine (5 min), fixed with 4% formaldehyde (15min) and permeabilized with Reduction Solution (10 min). Riboprobes (*Smed-pctk-1, Smed-sim, Smed-gpas, Smed-ovo1, Smed-tph, Smed-piwi1*) were synthesized using the DIG RNA labelling kit (Sp6/T7, Roche). Animals were mounted in 80% glycerol before imaging.

### Immunohistochemistry staining

Whole-mount immunohistochemistry was performed as previously described [35,72]. The primary antibody used were: mouse anti-VC1 [17], specific for planarian photosensitive cells (anti-arrestin, kindly provided by H. Orii and Professor K. Watanabe) diluted 1:15000 as well as rabbit anti-phospho-histone-3 (Ser10) to detect cells at the G2/M phase of cell cycle (H3P, Cell Signaling Technology) diluted 1:300. The secondary antibodies Alexa 568-conjugated goat anti-mouse and goat anti-rabbit (Molecular Probes, Waltham, MA, USA) were diluted 1:1000. Nuclei were stained with DAPI (1:5000; Sigma-Aldrich). Samples were mounted in 70% glycerol before imaging.

### Planarian phototaxis assays

Phototaxis assays were performed following a simplified protocol adapted from [18]. Planarian phototaxis responses were recorded over a 180-second period using an overhead digital video camera (Canon EOS550D). To generate a directional light gradient, the container was protected by a black screen with a hole that allowed the entrance of 500 lux of white light from one side of the container, as shown in Figure 6A.

### Microscopy, image processing and quantification

Live animals were imaged using an sCM EX-3 high end digital microscope camera (DC.3000s, Visual Inspection Technology). Fixed and stained animals were observed with a Leica MZ16F stereomicroscope and imaged with a ProgRes C3 camera (Jenoptik, Jena, TH, Germany). Image processing was performed with Adobe Photoshop 2024. Quantification of pigment cup area and total eye area (Fig. 2) was carried out manually and normalized by the area of the animals using ImageJ-win64. Quantification of *ovo1*+, *tph+* and VC1+ cells (Fig. 3) was carried out manually using ImageJ-win64. Quantification of the length of the photoreceptor axonal projections (Fig. 4) was carried out manually and normalized by the length of the brain or the animals using ImageJ-win64. Quantification of *sim*+ cells (Fig. 5) was carried out manually using ImageJ-win64. Quantification of distance covered in cm (Fig. 6) in the phototaxis assays was carried out manually measuring the initial and the final position of the animals.

### Phylogenetic analyses

Protein sequences of CDKs homologues were obtained from NCBI and Planmine v3.0 [73] and CDKs protein sequences from other organisms were obtained from [74] (supplementary material S1). All sequences were aligned using MAFFT with the G-INS-i strategy [75]. The resulting full-length alignment was trimmed and used for phylogenetic reconstruction. Phylogenetic analyses were performed with the IQ-TREE Web server using default settings, including automatic selection of the substitution model, ultrafast bootstrap analysis with 1000 replicates and single branch testing with 1000 replicates [76–77]. The approximate Bayes test option was selected. The phylogenetic tree was visualized using iTOL (https://itol.embl.de) and edited with Adobe Illustrator.

### Statistical analysis and graphical representation

To identify significant differences between conditions, unpaired Student’s t test was performed after first confirming data normality and homogeneity using the Shapiro-Wilk test. Error bars represent the standard error of the mean (SEM). All the analyses were done using GraphPad Prism.

## Supporting information

Guixeras_Supplementary

## Abbreviations

CDKs: cycline dependent kinases
CNS: central nervous system
DAPI: 4’,6-Diamidino-2-Phenylindole (DNA stain)
dR: days of regeneration
dpi: days post injection
dsRNA: double stranded RNA
FISH: Fluorescent *In Situ* Hybridization
PCTKs: PCTAIRE kinases
Pctk-1: PCTAIRE kinase 1 (CDK family subfamily)
PH3: Phospho-Histone H3 (mitosis marker)
Q1, Q2, Q3, F: Quadrants and final region used in phototaxis assays RNAi RNA interference
Smed: *Schmidtea mediterranea*
WISH: Whole-Mount *In Situ* Hybridization
X0, X1, Xf: Initial, intermediate, and final positions in movement assays

## Acknowledgements

We wish to thank all members of the Francesc Cebrià lab for their suggestions and discussion of the results.

## Author’s contributions

A.G-F. performed the experiments and analysis of the data. A.G. participated in the experiments. A.G-F., M.D.M and F.C contributed to the conception of the work, the interpretation of the data and drafting and revision of the manuscript. All authors read and approved the final manuscript.

## Funding

This work was supported by grants PID2021-126958NB-I00 (Ministerio de Ciencia, Innovación y Universidades, Spain) and 2021SGR00293 (AGAUR, Generalitat de Catalunya) to F.C, and grant PID2024-159121NB-I00 (Ministerio de Ciencia, Innovación y Universidades, Spain) to F.C and M.D.M. A.G-F. was supported by a FPU fellowship (Ministerio de Ciencia, Innovación y Universidades, Spain). M.D.M. has received funding from the programme Beatriu de Pinós, funded by the Secretary of Universities and Research (Government of Catalonia) and by the European Union Horizon 2020 research and innovation programme under Marie Sklodowska-Curie grant agreement No. 801370 (2018BP-00241).

### Data availability

All data generated or analyzed during this study is included in this manuscript and its additional files.

## Declarations

### Ethics approval and consent to participate

Not applicable

### Consent for publication Not applicable

