## Supplementary material for "A PCTAIRE family kinase regulates eye and brain size in freshwater planarians": Guixeras_Supplementary

### Supplementary Information

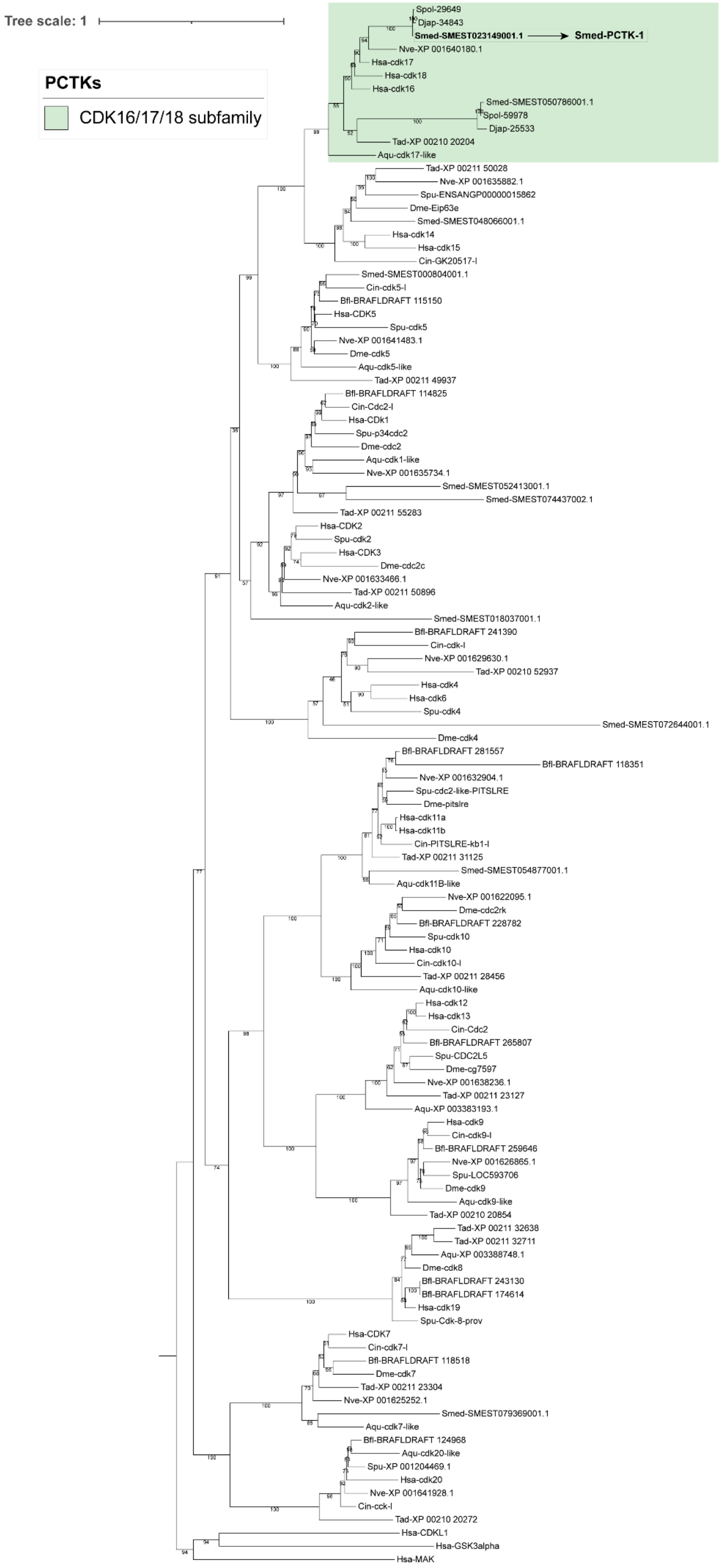

Additional file 1. Figure S1. Phylogenetic tree of the CDK family proteins**.** The phylogenetic relationships between Smed-CDKs and other metazoan CDK proteins were inferred using a maximum likelihood (ML) approach. The full-length aligned CDK sequences served as the basis for constructing the tree. Numbers shown above the branches represent ML bootstrap support values. Members of the PCTK subfamily are marked in green, with Smed-PCTK-1 specifically indicated. The scale bar reflects the number of substitutions per site. HsaGSK3α, HsaMAK, and HsaHCDKL1 were included as outgroup sequences. All proteins are annotated with their species names followed by their accession numbers. Species abbreviations are as follows: Smed, *S. mediterranea*; Spol, *S. polychroa*; Djap, *D. japonica*; Hsa, *H. sapiens*; Cin, *C. intestinalis*; Spu, *S. purpuratus*; Bfl, *B.a floridae*; Dme, *D. melanogaster*; Nve, *N. vectensis*; Tad, *T. adhaerens*; Aqe, *A. queenslandica* (original sequences extracted from Planmine v3.0 [73] and [74]).

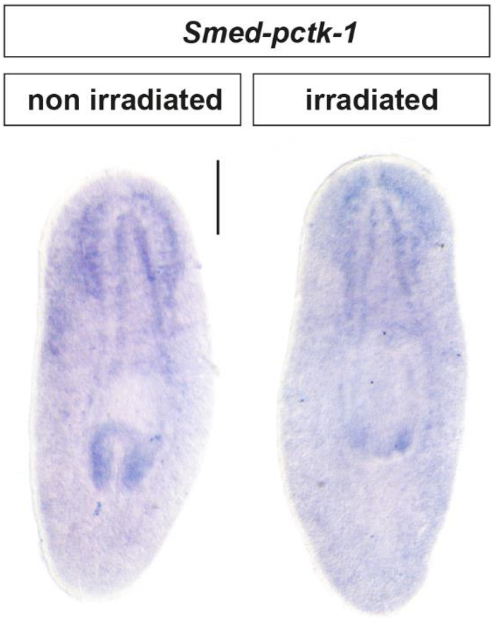


Additional file 2. Figure S2. *Smed-pctk-1* expression pattern in non-irradiated and irradiated intact animals detected by whole mount *in situ* hybridization (WISH). Scale bar: 400 µm.

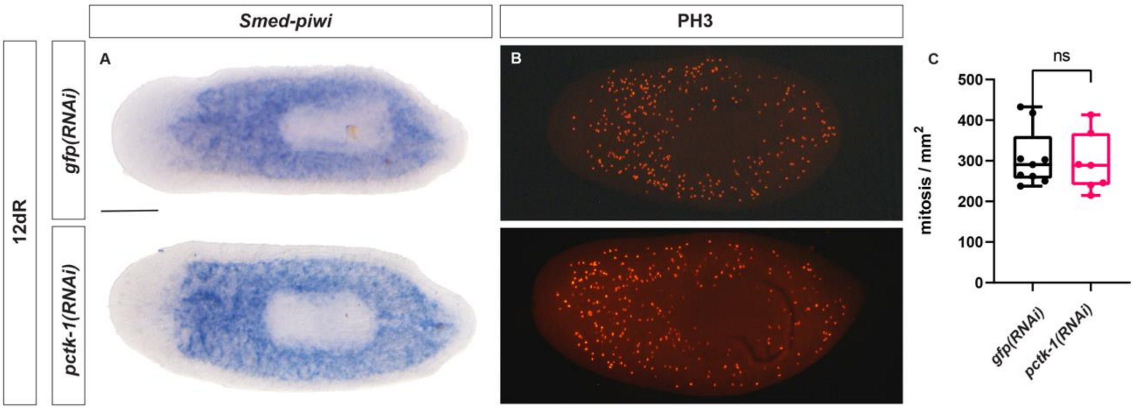


Additional file 3. Figure S3. *Smed-pctk-1* silencing does not affect the stem cell populations. **A** Neoblast population labelled by WISH for *Smed-piwi-1*. **B** Mitotic cells visualized by anti-PH3 immunostaining. **C** Number of mitotic cells per mm$^2$ ($^{ns}$p-value >

0.05, Student's t-test). Values represent the mean of at least 8 animals per condition. Animals at 12 days of regeneration (dR). Scale bar: 300 µm.

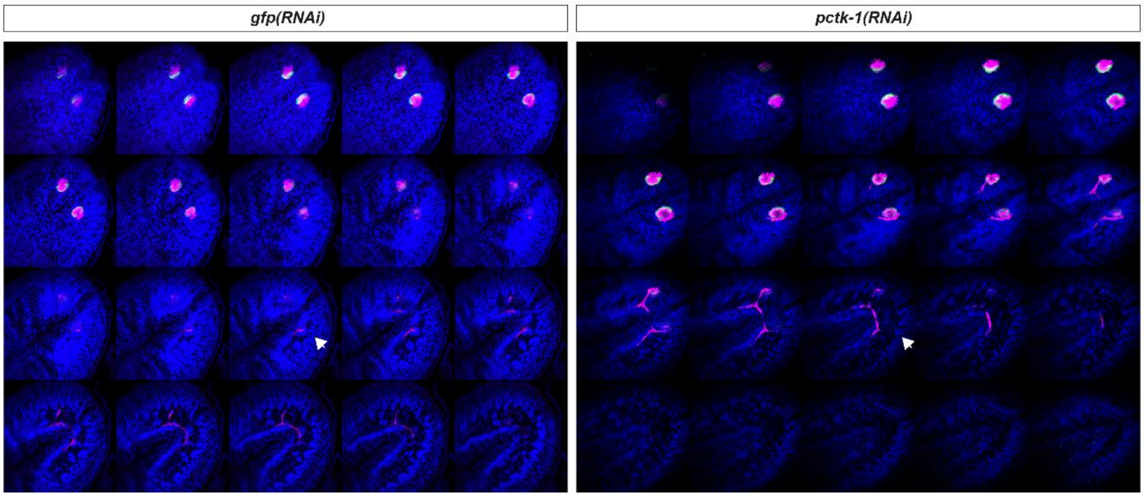


Additional file 4. Figure S4. Photoreceptor cells are abnormally present in more ventral locations. Optic chiasm and photoreceptor neurons visualized by VC-1 immunostaining (in magenta), eye pigment cells visualized by *tph* FISH (in cyan) and cell nuclei by DAPI (in blue). The images show consecutive confocal sections revealing that, in *Smed-pctk-1* silenced animals, photoreceptors cells reach more ventral regions closer to the cephalic ganglia (white arrowheads). Scale bars: 200 µm. Images at 53 days after amputation (dR).

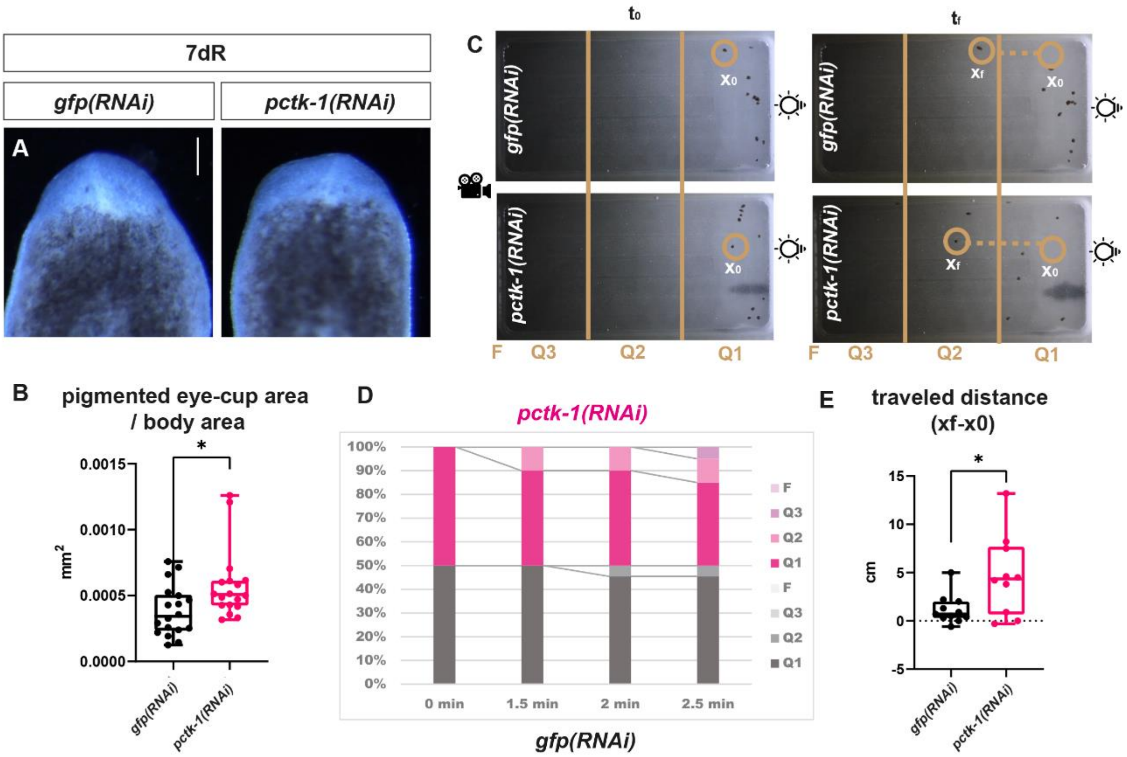


Additional file 5. Figure S5. Increased light sensitivity in tails after *Smed-pctk-1* silencing. **A** Live tails at 7 days of regeneration (dR). Scale bar: 300 µm. **B** Ratio between the pigmented eye-cup area and the total body area. (*p-value < 0.05, Student's t-test). Values represent the mean of at least 12 eyes per condition. **C** Schematic representation of the phototaxis behavioral assay. **D** Light responses measurements by quantifying the percentage of animals in Q1, Q2, Q3 and F regions at various time points after the exposition of direct light. **E** Light responses measurements by calculating the distance in cm (Xf-X0) after the exposition to direct light. (*p-value < 0.05, Student's t-test). Values represent the mean of at least 10 animals per condition. Animals are at 7 days of regeneration.
